# Improving the Efficiency of Genomic Selection in Chinese Simmental beef cattle

**DOI:** 10.1101/022673

**Authors:** Jiangwei Xia, Yang Wu, Huizhong Fang, Wengang Zhang, Yuxin Song, Lupei Zhang, Xue Gao, Yan Chen, Junya Li, Huijiang Gao

## Abstract

Genomic selection is an accurate and efficient method of estimating genetic merits by using high-density genome-wide single nucleotide polymorphisms (SNPs).In this study, we investigate an approach to increase the efficiency of genomic prediction by using genome-wide markers. The approach is a feature selection based on genomic best linear unbiased prediction (GBLUP),which is a statistical method used to predict breeding values using SNPs for selection in animal and plant breeding. The objective of this study is the choice of kinship matrix for genomic best linear unbiased prediction (GBLUP).The G-matrix is using the information of genome-wide dense markers. We compare three kinds of kinships based on different combinations of centring and scaling of marker genotypes. And find a suitable kinship approach that adjusts for the resource population of Chinese Simmental beef cattle. Single nucleotide polymorphism (SNPs) can be used to estimate kinship matrix and individual inbreeding coefficients more accurately. So in our research a genomic relationship matrix was developed for 1059 Chinese Simmental beef cattle using 640000 single nucleotide polymorphisms and breeding values were estimated using phenotypes about Carcass weight and Sirloin weight. The number of SNPs needed to accurately estimate a genomic relationship matrix was evaluated in this population. Another aim of this study was to optimize the selection of markers and determine the required number of SNPs for estimation of kinship in the Chinese Simmental beef cattle.

We find that the feature selection of GBLUP using Xu’s and the Astle and Balding’s kinships model performed similarly well, and were the best-performing methods in our study. Inbreeding and kinship matrix can be estimated with high accuracy using ≥12,000s in Chinese Simmental beef cattle.

## Introduction

With the development of genetic markers, especially high throughput genotyping technology, it becomes available to estimate breeding value at genome level, i.e. genomic selection (GS)[1,2]. Genomic selection increases the rate of genetic improvement and reduces cost of progeny testing by allowing breeders to preselect animals that inherited. Several approaches of genomic prediction have been presented. One of them is the genomic best linear unbiased prediction (GBLUP), which uses Genomic information in the form of a genomic relationship matrix that defines the additive genetic covariance between individuals[3,4]. The genomic relationship coefficients are estimated with higher accuracy than when using pedigree information because genomic information can capture of Mendelian sampling across the genome. GBLUP has become popular approach in genomic selection of dairy cattle [5,6], because it is simple and has low computational requirements[7,8].Traditionally relationships are encoded in pedigrees of known relatives[9-11], but for more distantly related individuals, pedigree information can sometimes be erroneous or difficult to obtain. Relatedness can also be calculated from large panels of genetic markers[12-16]. The other is to predict GEBV with genetic relationship matrix, which constructs genetic relationship matrix via high throughput genetic markers and then predicts GEBV through linear mixed model (GBLUP)[17]. Genomic predictions can be based on a BLUP-GS model in which the average relationship matrix based on pedigree in the traditional BLUP model is replaced by a genomic relationship matrix based on markers[18]. The expected relationship matrix among individuals in the population is replaced with the realized relationship matrix (or genomic relationship matrix) derived from markers[19].

At the same time new genotyping technologies have contributed to a reduction of genotyping costs. However, the cost of genotyping with a large number of markers is still a barrier for practical application of a G matrix in Chinese Simmental beef cattle breeding programs. A reduced set of markers which could estimate an accurate G matrix would contribute to substantial cost reduction of genomic selection schemes. Our research also find the number of markers we can get to estimate genomic inbreeding coefficient and kinship matrix accurately in Chinese Simmental beef cattle. To have a comparison, we also used a least absolute shrinkage and selection operator (LASSO) approach to estimate marker effects for genomic selection. Compare with the GBLUP approach to find their superiorities.

In this article, cross validation (CV) is applied to assess prediction ability[20,21]. In the cross validation method, the basic idea is to divide a data set into a training set and a validation set, to omit any kind of information of the validation set and to predict this information. However, in animal breeding applications individuals present vary degrees of genetic relationships, and obtaining independent training and validation sets is seldom possible.

Genomic relationships can better estimate the proportion of chromosomes segments shared by individuals because high-density genotyping identifies genes identical in state that may be shared through common ancestors not recorded in the pedigree. A genomic relationship matrix (G) can be calculated by different methods. We investigate three kinship matrices within genetic best linear unbiased prediction (GBLUP). And we find a suitable kinship approach that adjusts for the resource population of Chinese Simmental beef cattle. The another aim of this study was to optimize the selection of SNPs and determine the number of informative SNPs necessary to estimate genomic inbreeding and kinship matrix accurately in Chinese Simmental beef cattle.

## Materials and methods

### Ethics statement

The whole procedures we do for animals were in strict accordance with the guidelines proposed by the China Council on Animal care, and all protocols were approved by the Science Research Department of the Institute of Animal Science, Chinese Academy of Agricultural Sciences (CAAS) (Beijing, China). The use of animals and private land used in this study were approved by the owners. And samples were collected along with the regular quarantine inspection on the farms.

#### Animal resource and phenotypes

Our resource population of the Simmental cattle was established in Ulgai, Xilingol league, Inner Mongolia of China,the mapping population consisted of 1059young Simmental bulls born in 2009-2014. After weaning, the cattle was moved to Beijing Jinweifuren cattle farm for feedlot finishing under the same feeding and management system. Each individual bull was observed for growth and developmental traits until slaughtered at 16-18 months of age. This study mainly focused on the phenotypic traits associated with cattle meet production, so during the period of slaughter, carcass traits and meat traits were measured according to the Institutional Meat Purchase Specification for fresh beef guide lines. Two carcass related traits were studied in the study. Carcass weight (CW) was measured after slaughter and bloodletting by eliminating the hide, head, feet, tail, entrails and gut fill. And sirloin weight(SW) were measured directly from carcass anatomy.

#### Phenotypic correction

After collecting the original data, phenotypes should be correctedin advance. With the fixed effects, including years, farms and fatten daysof birthentering weight using the following equation:

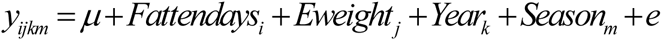

Where *y*_*ijkm*_ is the phenotypic value, **μ** is the population mean, Both of *Fattendays*_*i*_ and *Eweight*_*j*_ are continue variables, *Fattendays* is the days since entering fattening farm to slaughtering and *Eweight*_*j*_ is the live weight when entering fattening farm. *Year*_*k*_ is the slaughtering year, which are divided into three groups (2009, 2010 and 2011, 2012, 2013, 2014). *Season*_*m*_ is the calving season including the three levels (November to April, May to August and September to October). e was the random residual. which was exerted for the subsequent association study with SNP.

#### SNP data

Semen or blood samples were collected along with regular quarantine inspection of the farms. Genomic DNA was extracted from blood samples using a TIANamp Blood DNA Kit (Tiangen Biotech Company limited, Beijing, Chain), and DNAs with an A260/280 ratio ranging between 1.8 and 2.0 were subject to further analysis. All individuals were genotyped using the Illumina BovineSNP BeadChip containing 774660 SNPs,

#### Genotyping and quality control

Blood samples we collected along with regular quarantine inspection of the farms. Genomic DNA was extracted from blood samples using a TIANamp Blood DNA Kit (Tiangen Biotech Company limited, Beijing, Chain), and DNAs with an A260/280 ratio ranging between 1.8 and 2.0 were subject to further analysis. the mean value of distance between each marker is 3.43Kb and variance value of distance between each marker is 19.19Mb The Illumina BovineHD BeadChip[22] contains 774,660 SNPs were manufactured for individuals genotyping.

About quality control. We used The PLINK software (v1.9, http://pngu.mgh.harvard.edu/∼purcell/plink/) to exclude individuals and remove undesired SNPs. The procedure about quality control was conducted as follows: firstly, when call rates are less than 90%, minor allele frequencies are less than 5%, genotype appearances are less than five individuals or departure from Hardy-Weinberg equilibrium is severe (with lower than 10^−6^ probability). Then an individual would be excluded with missing genotypes above 10% or Mendelian error of SNP genotype more than 2%. Additionally, all of the misplaced SNPs were excluded from the analysis.

#### Model

We explored the effects of the GBLUP with different kinship matrixes on the predictive power of GS models using Chinese beef cattle real data sets including continuous phenotypic traits about Carcass weight and sirloin weight.

Mixed model for GBLUP

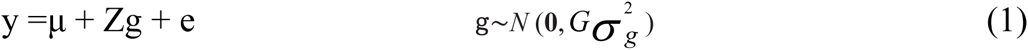

where g are the random effects and Z is a design matrix that can be used for example to indicate the same genotype exposed to different environments. Any positive definite matrix can be used for G. Fixed effects can also be included in (1) in order to capture purely environmental effects. where G is the realized genomic relationship matrix, calculated from marker genotypes without using pedigree information. Following VanRaden[18].

#### Kinship Estimation

Genomic relationships can better estimate the proportion of chromosomes segments shared by individuals because high-density genotyping identifies genes identical in state that may be shared through common ancestors not recorded in the pedigree. The SNP-based kinship of two individuals is usually based on the average over SNPs of the product of their genotypes, coded as 0, 1 and 2 according to the count of one of the two alleles. By design, it can only capture the additive components of kinship, and it has very low power in identifying non-additive ones. In the following, we denote this genotype matrix with **X**, with rows corresponding to individuals and columns to SNPs, and with *x*_*i*_ its *ith* column.

The first study that presented a marker based relationship matrix was VanRaden et al., [18]. They calculated the relationship based on the concept of a similarity index. When the marker data have been collected on a sample of individuals, we can estimate the VanRaden’s kinship matrix[18,23] as

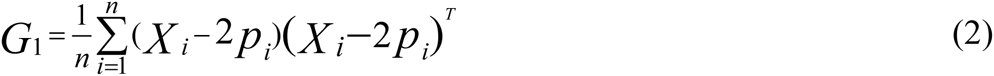

In this kinship matrix.**n** is the number of markers and _*pi*_ is a vector with every entry equal to the population allele fraction. Centring improves interpretability, since kinship values can be interpreted as an excess or deficiency of allele sharing compared with random allocation of alleles. and so zero can be interpreted as “unrelated”. However, the requirement to estimate the _*pi*_, usually from the same data set, can cause problems in some settings.

One criticism of the kinship matrix is that the sharing of a rare allele between two individuals counts the same as the sharing of a common allele. On enatural approach to giving more weight to the sharing of a rare allele is to standardize over SNPs. So we can estimate the Astle and Balding’s kinship matrix[16,24]as

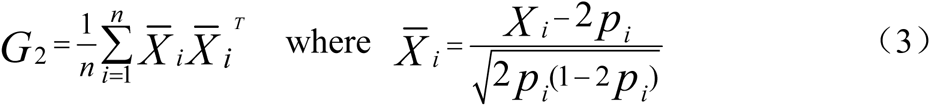

The (i, j) entry of *G*_2_ can be interpreted as an average over SNPs of the correlation coefficient estimated from a single pair of individuals, i and j.

For the *k*th marker from the *j*th animal, the SNP genotype was numerically coded as *X _jk_* = 1, *X _jk_* = 0 and *X _jk_* = −1, respectively, for the homozygote of minor allele, the heterozygote and the homozygote of major allele, *X _jj_* is the situation of *j* equal *k*. Therefore, the genotype data are represented by a *Z* matrix with a dimensionality *n* × *m*, where *m* = 1059 the sample is size and *n* = 640000 is the number of SNP markers. The marker generated Xu’s kinship matrix[25] calculated using

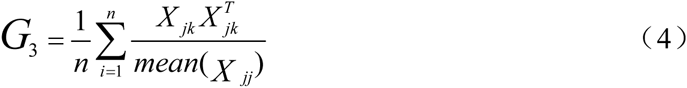

#### Least absolute shrinkage and selection operator (LASSO)

LASSO (citation) is a variable selection approach in statistics, but it has been used for genomic prediction (citation). Here, we adopted this method as the standard for comparison. Although Bayes B (citation) is a more desirable standard method for comparison, it may take unreasonable amount of time to finish the data analysis for such a large number of markers in the data.

#### Cross validation

Cross validation (CV)[20,21], sometimes called rotation estimation, is a technique for assessing how the results of a statistical analysis will generalize to an independent data set. It is mainly used in settings where the goal is prediction, and one wants to estimate how accurately a predictive model will perform in practice. In k-fold cross-validation, the original sample is randomly partitioned into k equal size subsamples.

In this study, we used five types (2,4,6,8,10) of cross validation to analysis the impacts of varying the training set size on the predictive ability. Each type of CV was replicated 200 times, resulting 200 average predictive abilities. In one replicate of a CV, the entire set is randomly divided into a training set, which is used for parameter estimation; and a validation set, for which genetic values are predicted. The predictive abilities are then averaged to obtain one average correlation per CV replicate.

#### Construction of Marker Panel Subsets

The R software was used to test the number of markers necessary to precisely predicted breeding values withthe approach of GBLUP in Chinese Simmental beef cattle. Subsets (n= 600, 1500, 3000, 6000, 12000, 3000, 60000, 120000, 240000, 360000, 680000.) of markers were randomly sampled with replacement from the full set of 680000 markers to ensure a random representation of the entire genome within the marker subset. We choose equal markers from every chromosome(30) for all 1059 cattle. And using the subsets of markers to estimate the kinship matrix.

## Results

### Effect of predictive correlations with different number of markers

The first we use different marker sets (n= 600, 1500, 3000, 6000, 12000, 3000, 60000, 120000, 240000, 360000, 680000.) to estimate kinship matrix. And then from the GBLUP we can get the predictive correlations10-fold _*pcv*_ (i.e. the correlations obtained from cross-validation), cross validation (CV) is applied to assess prediction ability with different number of markers. As can be seen from Figure S1. While estimates of genomic relationship matrix based upon at least 12,000 SNPs appear to be extremely stable, the predictive correlations can be robust. Estimating appear to be very sensitive to SNP sample size when fewer than 6,000 SNP are used. It has very significant consequences for both conservation genetic and GS applications because there are currently a question about cost of the high throughput sequencing.

The results suggest that the small sets of SNPs are also likely to be commercialized within the beef and dairy cattle industries. It will have some utility for the estimation of genomic relationship matrix and that will allow the estimation of molecular breeding values for traits other than those targeted by the SNPs within the panels. However, our results also indicate that the greatest benefits of the technology will not be realized until inexpensive assays can be produced which query ≥12000 SNPs.

### Choose a suitable kinship matrix estimation for Chinese Simmental beef cattle

We evaluated predictive ability for CW and SW using three kinds of kinship matrix approaches via a series of 2,4,6,8,10-fold CVs with all markers and get 200 replicates. From figure S2 and S3 we see that the predictive performance of GBLUP improves as the kinship matrices progress from G1 through to G3. G2.The results are illustrated in table. S1. The means of the predicted correlations of G3 in the 2 4 8-fold are higher than G2 on the trait of CW, and they are both more accuracy than G1. While in the trait of SW from table. S2, the predicted correlations of 4 6 8 10-fold are higher than G3, and they both have a better effect than the G1. Although small differences were observed in the ranks obtained with different genomic matrices. However, these differences have direct implications on selection decisions and genetic progress.

### Comparison with the method of LASSO

Another aim of this study was to compare the predictive abilities for CW and SW using GBLUP method and the LASSO method. And we can compare them to find their superiorities. In our study, the candidate population GEBV accuracy with the all markers was compared with using CV-GBLUP and CV-LASSO approaches (Table S3) via a series of 10-fold CVs with 100 replicates and 20 replicates. The accuracy obtained by GBLUP exceeded that of LASSO. Table 3 also shows the computing time needed to run LASSO was more than for GBLUP. A large proportion of the time in LASSO is spent in cross-validation used to define the markers size.

## Discussion and conclusions

Simmental beef producers select for meat yield and quality to increase their income from steer feedlots and sold sirloin. Estimated breeding values (EBVs) for Carcass weight (CW), sirloin weight(SW) are commonly used as selection criteria in attempts to increase meat yield and quality, which determine profitability for the Simmental beef industry[26].To our knowledge, this is the first application of genomic prediction on a real set of SNP genotyping data in Chinese beef Simmental cattle. Moreover we have used two approaches of cv-LASSO and cv-GBLUP (with three kinship matrix methods) to obtain it.

Due to the huge number of SNPs, it is interesting to compare the accuracy with different number of markers to construct the kinship matrices. SNP density is an important factor affecting accuracy of prediction in previous papers[27].Based on the result of our real Simmental data analysis, the predictability was almost constant until the number of markers reduced to about 12000, below which the curve started to drop rapidly. That is to say we could get a stable estimation of kinship with no less than 12000 markers. In the current study it was shown that 2,500-10,000 SNPs were needed for robust estimation of genomic relationship matrices with high accuracy in cattle[28,29]. Furthermore 10,000 SNPs the genomic relationship coefficients seemed to be extremely robust while building the G matrix with 2,500 SNPs seemed to be very sensitive to SNP sample size. The inappropriate selection of marker numbers is a consequence of our previous real data result and other reports. They all provide the evidence that little change of predictive accuracy will occur when the number of markers used to construct the kinship is over 10,000. In another way some studies [30,31] found that little decrease in accuracy was obtained when the selected markers are low in useful LD with causal polymorphisms. The underlying mechanism therefore seems to depend on a sufficient number of SNPs being in low LD with causal polymorphisms, rather than few SNPs in close physical association and high LD. The present study confirmed that dcreasing the number of markers didn’ t result in reduction of information. Thus, using a reduced set of markers is possible to estimate accurate genomic inbreeding and kinship matrix estimation and makes it possible to reduce genotyping costs as well.

Second, the objective of this study was to apply different genomic matrices to analyses of high density SNP panel in a Simmental population and evaluate the impact of those G matrix on predictive correlations. It is obvious that different methods used to construct the kinship matrix may result in a totally different results. From the previous paper we find three methods of kinship matrix estimation. The first empirical formula proposed by VanRaden is widely accepted that it could efficiently reflect the genetic relationship[18].But through the results, we know that the accuracy of predictive correlations is not quite good for Chinese Simmental beef cattle population. So the VanRaden kinship estimation approach isn’t suitable for the population compare with the other two methods. The Astle and Balding’s model is another widely adopted method to estimate the kinship matrix with markers[24].The first reason why we choose the method is the use of Astle and Balding’s model instead of identityby-state (IBS) model (another approach) slightly increased the observed reliability of the predictions. The second theAstle and Balding’s model often used in GWAS research and proved it had a very good effect[32-34]. Compared with the two models, Xu’s model also can provide the close relationship with markers. We first attempt to introduce this method for our research. At last we find this method also has an equal function with the Astle and Balding’s method. So the feature selection of GBLUP using the last two kinships performed similarly wellon Chinese Simmental beef cattle.

We also compared the performance of two different statistical methods to ensure that there are no artifacts caused by human errors in any single method. Two methods produced very similar results. Our results demonstrate that GBLUP can accurately estimate the effects of SNPs associated with QTL in dense SNP data, leading to accurately predicative correlation for genomic selection, in GBLUP, all SNP effects are assumed to be distributed normally, and the effects are fitted using a single distribution. But in LASSO, major SNPs can be selected from high-throughput SNPs, in this method, most SNPs are shrunk to zero and only a few key SNPs are retained, and thus LASSO can be used for selecting potential candidate SNPs. But in our results, we also find the method of cv-LASSO isn’t stable, the result prove it slightly floating, on the other hand the LASSO spend more time than GBLUP.

## Acknowledgements

This work was supported by the Cattle Breeding Innovative Research Team (cxgc-ias-03), the 12th “Five-Year” National Science and Technology Support Project (2011BAD28B04) basic research fund program, Chinese Academy of Agricultural Sciences Fundamental Research Budget Increment Project (2013ZL031), National High Technology Research and Development Program of China (863 Program 2013AA102505-4) and National Natural Science Foundations of China (31372294).

## Authors’ contributions

HJG and JYL conceived and designed the experiments. JWX and YW performed the experiments. HZF, WGZ, XYS analyzed the data. XG, YC and LPZ participated in the experiment. HJG and JYL supervised the experiment. JWX wrote the paper. All authors read and approved the final manuscript.

---

**Fig. 1.**
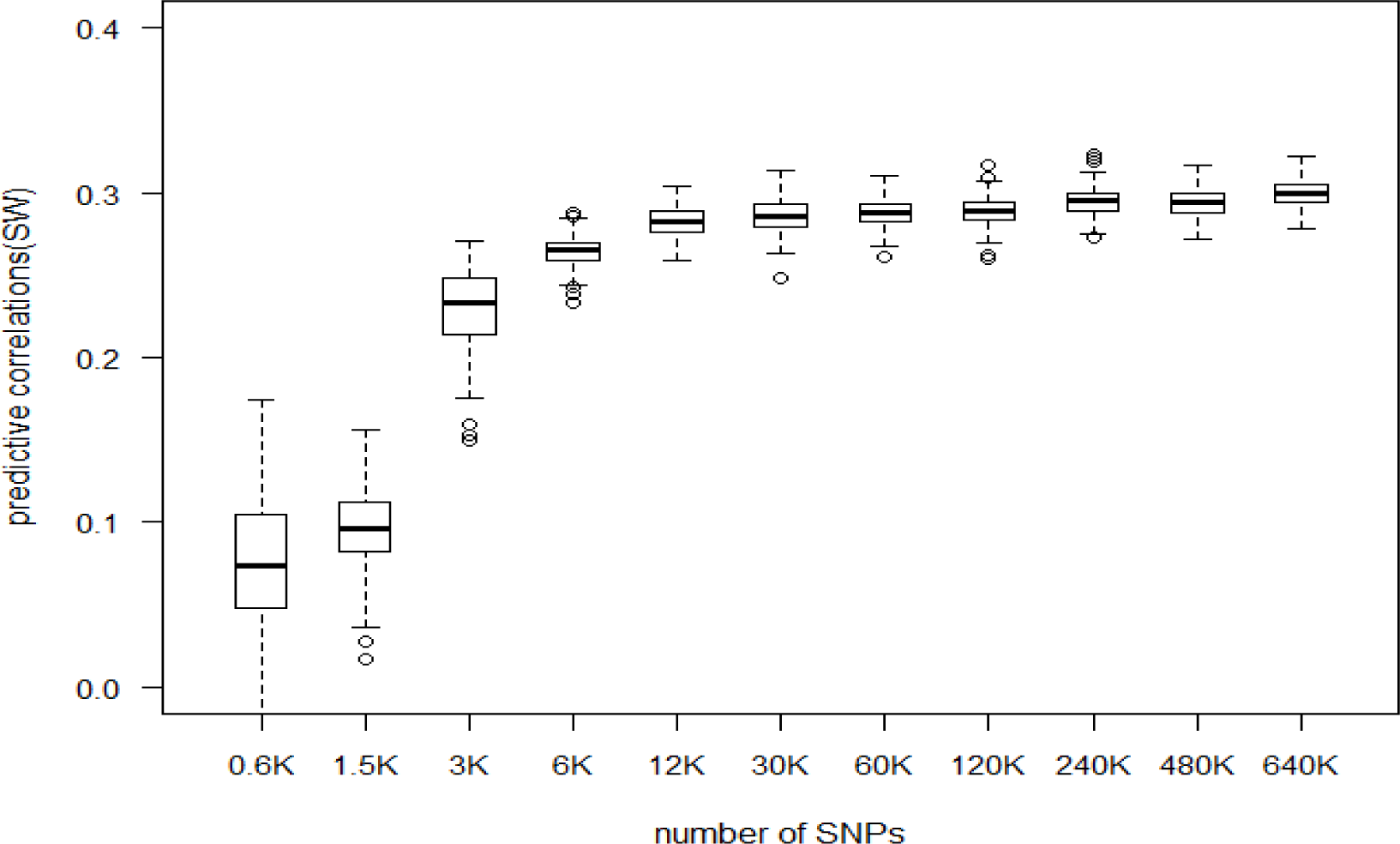
The predicted correlations with different SNP densities. The GBLUP of relationship matrix methods G2 we can get the 10-fold with 200 replicates to assess predicted correlations with different number of markers on the trait of sirloin weight (SW).

**Fig. 2.**
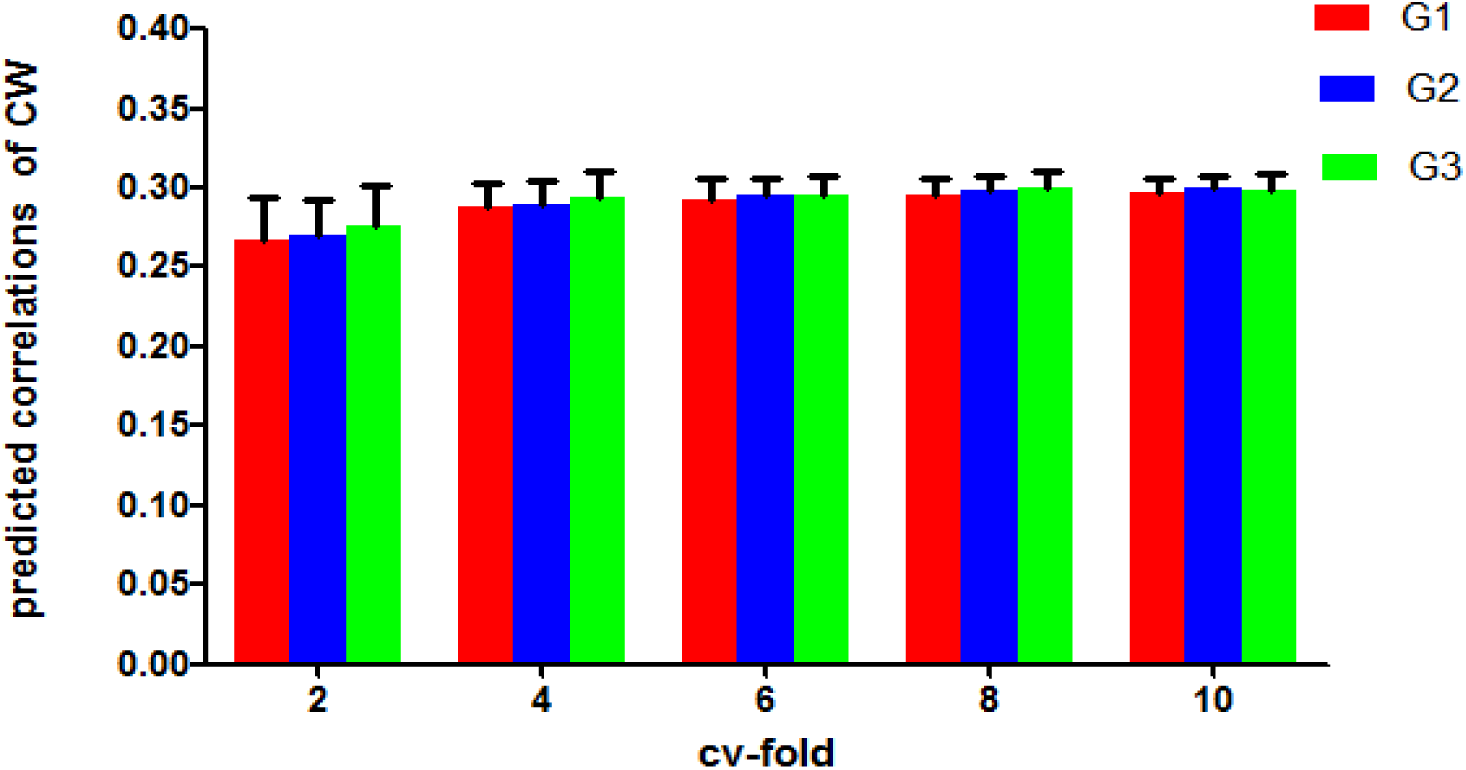
The predicted correlations of GBLUP with 2, 4, 6, 8, 10 cv-fold of CW using G1, G2, G3. Each Column chart illustrates the average predictive correlations for 200 replicates of CV procedure using GBLUP.

**Fig. 3.**
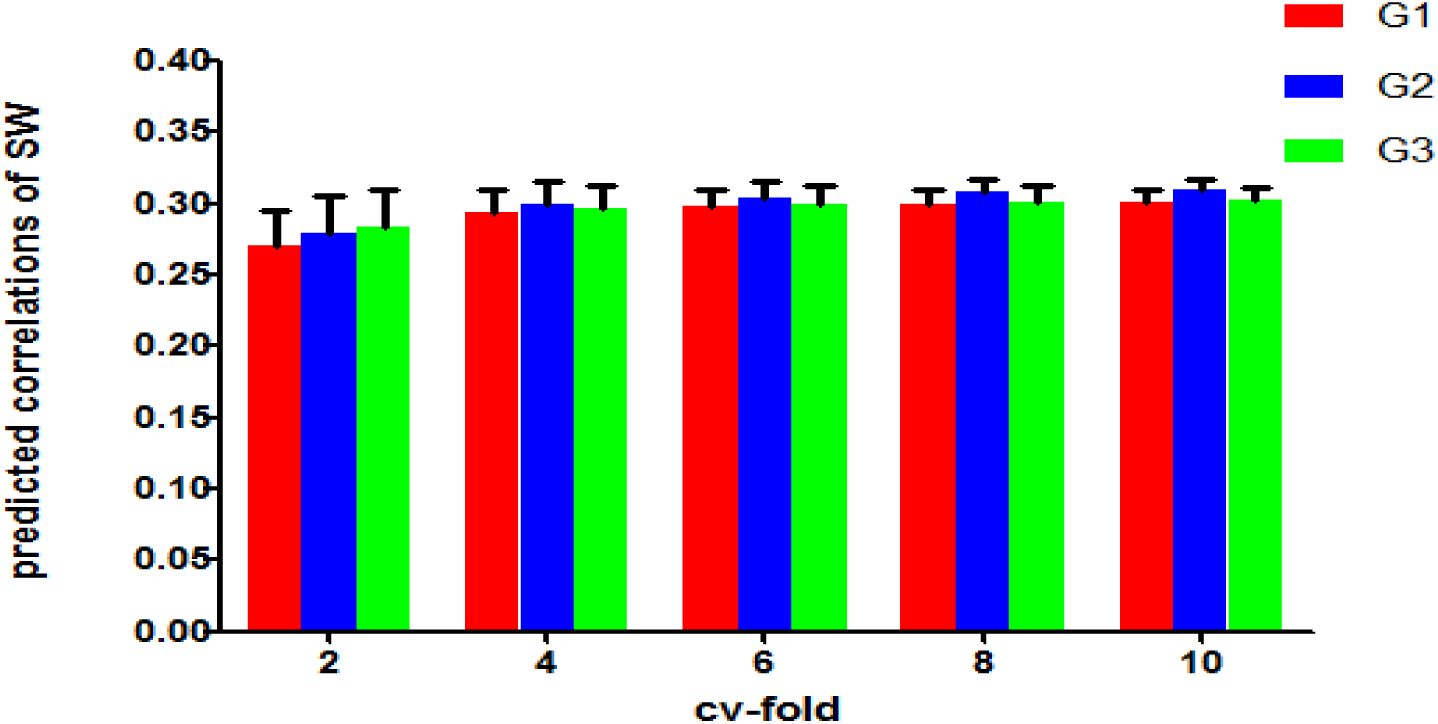
predicted correlations of GBLUP with 2, 4, 6, 8, 10 cv-fold of SW using G1, G2, G3. Each Column chart illustrates the average predictive correlations for 200 replicates of CV procedure using GBLUP.

**Table 1.** The mean of predicted correlations and with the Standard Deviations of GBLUP with 2, 4, 6, 8, 10 cv-fold of CW using G1, G2, G3.

| CV | G1(sd) | G2(sd) | G3(sd) |
| --- | --- | --- | --- |
| 2 | 0.2673(0.0271) | 0.2698(0.0232) | 0.2757(0.0255) |
| 4 | 0.2880(0.0158) | 0.2898(0.0143) | 0.2939(0.0163) |
| 6 | 0.2933(0.0122) | 0.2957(0.0105) | 0.2950(0.0122) |
| 8 | 0.2963(0.0102) | 0.2983(0.0096) | 0.2994(0.0108) |
| 10 | 0.2968(0.0095) | 0.2997(0.0082) | 0.2989(0.0100) |

**Table 2.** The mean of predicted correlations and with the Standard Deviations of GBLUP with 2, 4, 6, 8, 10 cv-fold of SW using G1, G2, G3.

| CV | G1(sd) | G2(sd) | G3(sd) |
| --- | --- | --- | --- |
| 2 | 0.2708(0.0250) | 0.2799(0.0251) | 0.2833(0.0261) |
| 4 | 0.2945(0.0147) | 0.2994(0.0157) | 0.2969(0.0155) |
| 6 | 0.2982(0.0113) | 0.3045(0.0105) | 0.3000(0.0133) |
| 8 | 0.2997(0.0107) | 0.3082(0.0089) | 0.3018(0.0104) |
| 10 | 0.3010(0.0083) | 0.3094(0.0084) | 0.3020(0.0090) |

**Table 3.** Predicted correlations, total computational time and memory resources required for the two methods used. They were calculated at 10-fold CV procedure for the SW trait with the similar computer, and get 200 replicates with GBLUP, 20 replicates with LASSO.

| method | CW | SW | Computing time<br>(min) | Allocated memory<br>(kb) |
| --- | --- | --- | --- | --- |
| GBLUP | 0.2997(0.0082) | 0.3094(0.0084) | 00:08:39 | 46368326 |
| LASSO | 0.2970(0.0427) | 0.2953(0.0412) | 02:03:29 | 44352028 |

**Table 3. Accuracy of prediction, total computational time and memory resources required for the two methods used.**
| method | CW | SW | Computing time<br>(min) | Allocated memory<br>(kb) |
| --- | --- | --- | --- | --- |
| GBLUP | 0.2997(0.0082) | 0.3094(0.0084) | 00:08:39 | 46368326 |
| LASSO | 0.2970(0.0427) | 0.2953(0.0412) | 02:03:29 | 44352028 |

